# A computational study on the optimization of transcranial temporal interfering stimulation with high-definition electrode using unsupervised neural network

**DOI:** 10.1101/2021.11.14.467844

**Authors:** Sang-kyu Bahn, Bo-Yeong Kang, Chany Lee

## Abstract

Transcranial temporal interfering stimulation (tTIS) can focally stimulate deep parts of the brain, which are related to specific functions, by using beats at two high AC frequencies that do not affect the human brain. However, it has limitations in terms of calculation time and precision for optimization because of its complexity and non-linearity. We aimed to propose a method using an unsupervised neural network (USNN) for tTIS to optimize quickly the interfering current value of high-definition electrodes, which can finely stimulate the deep part of the brain, and analyze the performance and characteristics of tTIS. A computational study was conducted using 16 realistic head models. This method generated the strongest stimulation on the target, even when targeting deep areas or multi-target stimulation. The tTIS was robust with target depth compared with transcranial alternating current stimulation, and mis-stimulation could be reduced compared with the case of using two-pair inferential stimulation. Optimization of a target could be performed in 3 min. By proposing the USNN for tTIS, we showed that the electrode currents of tTIS can be optimized quickly and accurately, and the possibility of stimulating the deep part of the brain precisely with transcranial electrical stimulation was confirmed.

## Introduction

Transcranial electric stimulation (tES), a method of stimulating the brain in a non-invasive approach by applying electric current to the scalp, has been attracting attention because of its safety^1^. This method is effective in enhancing brain cognitive function or treating psychopathy as an alternative to pharmacology^2–5^. Incipient tES attaches a large sponge near the area to be stimulated^6^. However, numerical analysis methods, such as the finite element method and boundary element method, are performed to analyze the electric fields of the brain. Thus, many electrodes, such as the international 10-10 system, can be used to stimulate the desired area more precisely^7, 8^. In addition to transcranial direct current stimulation (tDCS), which forms electric fields in direct current, the transcranial alternating current stimulation (tACS) method, which oscillates specific areas of the brain through alternating current stimuli, has also been used for electrical brain stimulation^9, 10^.

However, tDCS and tACS, which inject current to the scalp, have difficulty in stimulating the deep region because a large electric field tends to form around the electrodes^11, 12^. As an alternative, a method of obtaining only a mediating effect by stimulating the shallow region, which is speculated to be connected to the deep region, is used, rather than directly stimulating the deep place^13^. To overcome this, transcranial temporal interfering stimulation (tTIS), which uses the beat caused by interference of two different frequencies, has been proposed to modulate the deep region focally^14^. Numerous studies have been conducted on biological analysis and the phenomenon in the brain of the tTIS method^15–20^.

In contrast to tDCS and tACS, which enable linear combinations, tTIS is a nonlinear expression, and optimization of the electrode currents is challenging. In many studies, by comparing a number of combinations^21^ or adjusting the location gradually^20^, optimization is performed using a limited number of electrode sets, such as two pairs or more^20–25^. However, there are computational complexities to analyze and many limitations to stimulate precisely owing to the large number of electrode combinations.

An attempt to consider many electrodes has been made to optimize deep stimulation by using additional neck part electrodes in the international 10-10 system^26^. This is a method of optimizing in a sequential quadratic programming method^27^, with many constraints on the solution space consisting of each electrode current. This method continuously adjusts the power conditions to concentrate the stimulus. However, it is computationally complex and requires 20 to 40 hours to optimize one place.

The proposed method to optimize the electrode currents of tTIS uses unsupervised neural networks (USNNs). Neural networks are an approach to solve problems by imitating the connections between neurons in the human brain^28, 29^. Recently, this method has received considerable attention in various fields, with advances in hardware such as graphics processing unit computation^30, 31^ and massive amounts of databases^32^.

The mainstream neural network adopts the supervised training using the ground truth. However, the problem of finding electrode currents that stimulate the deep brain could not determine the correct answer set. Therefore, in this problem, a USNN is required. The main idea of this approach is to fix the specific formulas in the neural network of the feed-forward structure. The neural network is trained in a direction that minimizes the output of the network when an input is provided without a dataset. In this process, a specific connection weight corresponding to the actual output can be obtained. This approach has been widely used in nonlinear problems and in solving differential equations^33–37^.

In this paper, we aimed to propose a method using USNN to optimize the current of electrodes for tTIS. The basic concept of this method is that a neural network optimizes the set of electrode currents, compared with the target stimulus, using a fixed network converted from a stimulus formula. The proposed method performed optimizations using the electrode set of high-definition (HD) system faster than conventional studies and showed good performance in tests that stimulated two regions

## Material and method

### Realistic head model

The head models used in the simulation are “ICBM” data^38^ and averaged pediatric data^39, 40^, which are open data of brain magnetic resonance imaging images. These magnetic resonance imaging data were reconstructed to a five-layer finite element model consisting of the skin, skull, cerebrospinal fluid, gray matter, and white matter by using SIMNIBS^41–43^ (Fig.1a). A total of 69 electrodes were used for stimulation based on the international 10-10 system with a diameter of 1 cm. To attach these to the head, as shown in Fig.1b, the electrode was placed on an elliptical hemisphere similarly overlapped with the head and then projected to the nearest position of the head. Based on the anatomical information that cortical pyramidal neurons extend in the perpendicular direction^44^, the normal direction of the brain surface was used as the stimulation target. In particular, the interface between white matter and gray matter with a relatively good distinction between the elements was targeted. The normal direction of each surface node was determined by regressing one plane together with the adjacent surface node (Fig.1c). In this electrode system, the positions of all electrodes were assumed to produce two types of frequencies^26^.

**Figure 1.**
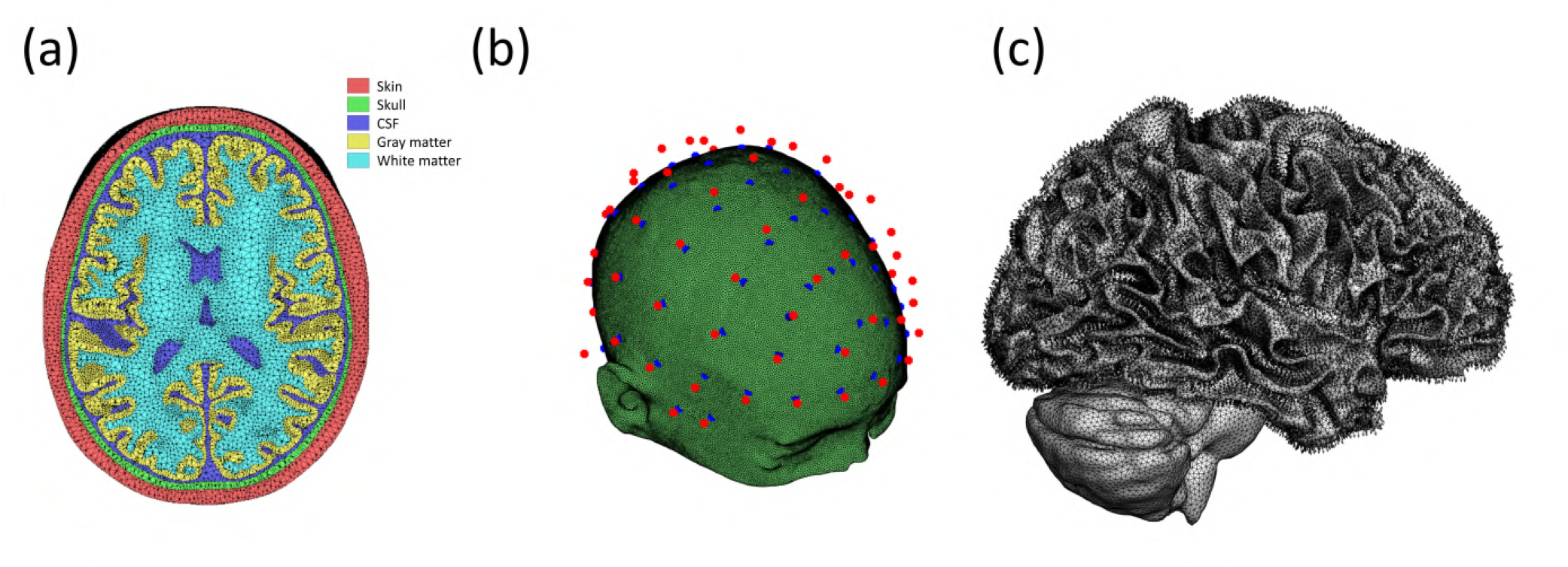
(a) Five-layer tetrahedral meshed finite element model consisting of the skin, skull, cerebrospinal fluid, gray matter, and white matter. (b) Electrode placement. Red dots indicate the positions placed based on the 10-10 system on the ellipsoid, and blue dots are the positions projected to the nearest scalp. (c) Normal unit vector plots on all nodes at the interface of white and gray matter.

### Mathematical formulation of stimulation

To calculate the electric potential V generated by the injected current through the electrodes, the Laplace equation ∇ · (*σV*) = 0 (*σ* : *electricconductivity*) was solved using the finite element method with linear approximation^45^. We could then obtain the electric field, which is defined as the negative gradient of the electric potential. The total electric field on the cortical surface, defined as the interfacial boundary between gray and white matter in this study, is represented in the following matrix form^46^:

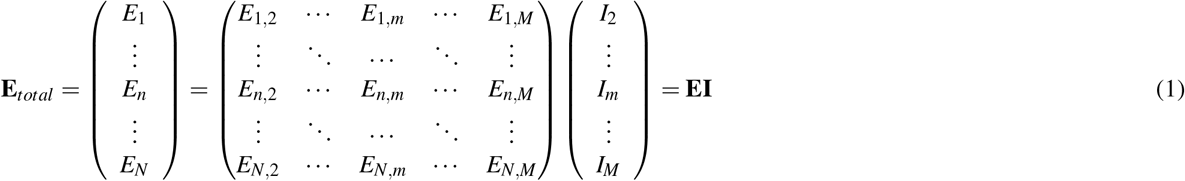

where, *I*_*m*_ is the electric current between the *m* – *th* electrode and the reference electrode, *E*_*n,m*_ is the electric field at the *n* – *th* node generated by the unit current between the *m* – *th* electrode and the reference electrode, and *E*_*n*_ is the total electric field at the *n* – *th* node. In the study, we considered the normal component of the electric field on the cortical surface, and the reference electrode was the first electrode.

### Modulation of tTIS

When two different high-frequency currents are injected on both sides of the head, a beat with an interference frequency corresponding to the difference between the two frequencies is generated. The frequency of the injected current is that the human brain cannot recognize and the difference between the two frequencies is the target frequency, and the modulation^26^ of the beat *Mod* is stronger in the deep region than in the shallow region. The equation for beat modulation is given by

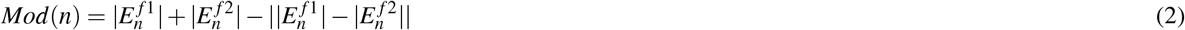

where 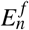 is the peak value for the electric field of the alternating current stimulation of frequency *f* at location *n*. Because the expression of beat modulation is nonlinear, the least square estimation cannot be applied to solve it, and a different approach for optimization is required.

### Structure of USNN

Figure 2a shows the tTIS optimization architecture. In this unsupervised training network, no input data are given, and unit constant 1 replaces this position. The nodes passing through the layers from unit constant 1 are fully connected to the electrode layer.

**Figure 2.**
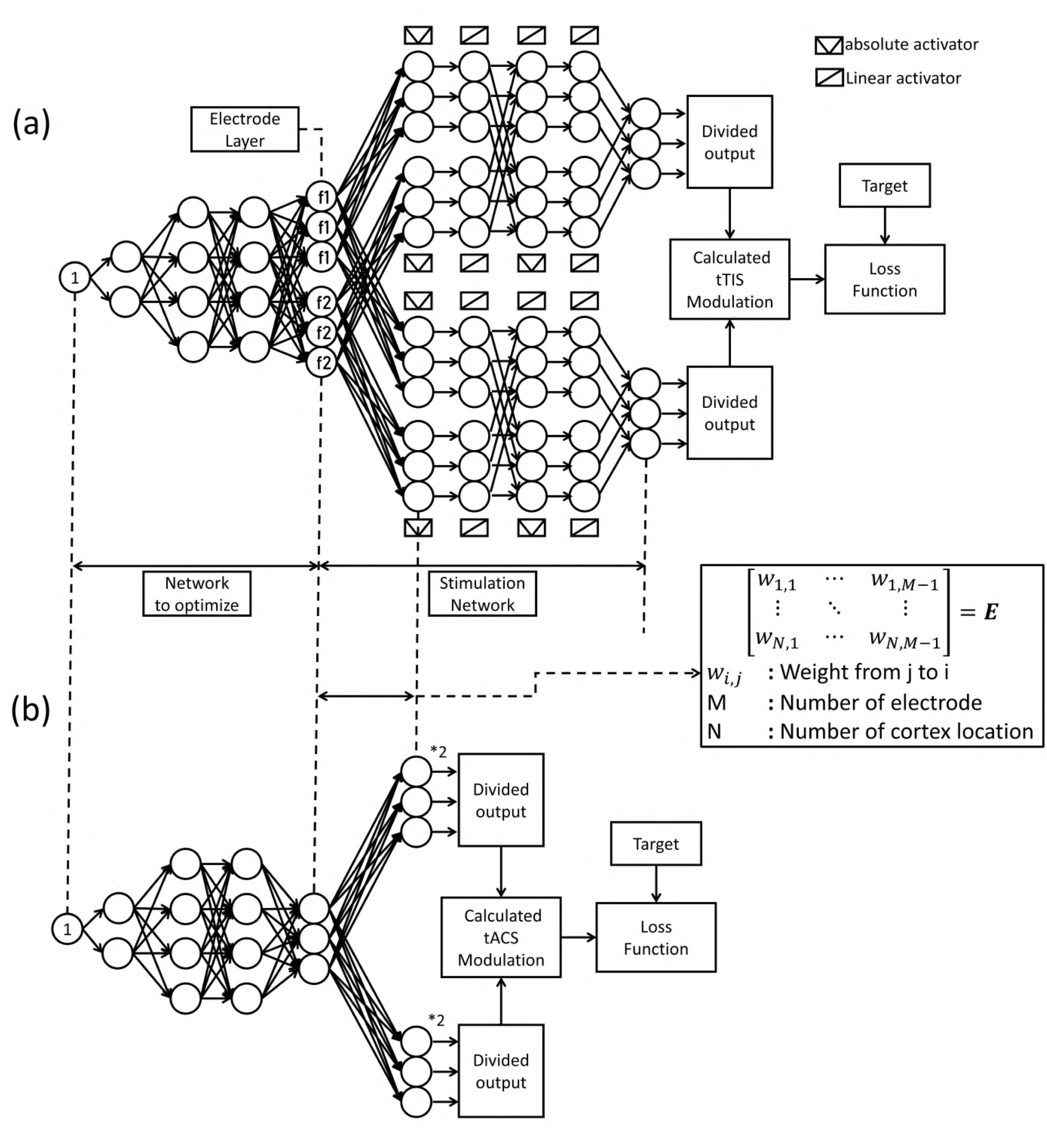
Architecture of USNN for electrode current optimization of (a) tTIS and (b) tACS.

The electrode layer has twice the number of electrodes used to use two different frequencies. The goal is to optimize the electrode currents that stimulate the target position focally by training these weights. Each node is connected by a ReLU activation function^47^, and after all the hidden layers, layer normalization^48^ is inserted to reduce the influence of the initial value and ensure stable convergence. Next of electrode layer, the stimulation network is connected to determine the modulation of the interference stimulus from the electrode nodes.

As described above, the electric field in the head can be obtained by a linear combination of the reference electric field and the currents for each electrode, as shown in Equation (1). Therefore, the fully connected connection between electrode nodes of a specific frequency and cortex location nodes is set as matrix **E** of Equation (1). Using these connections, the electric field of each frequency can be generated. Because a large number of locations exist in the cortex, they can be divided into appropriate sizes. To obtain the modulation of tTIS from the electric field amplitude for each frequency, a custom activation function that returns the absolute value of the input is declared and the electric field layers are connected as given in the formula in sequence. The entire network that calculates the modulation from the electrode has a fixed weight value that cannot be trained. The obtained modulation is compared with the given target and back-propagated to train the hidden layers to find the electrode currents. In the case of tACS, using a single frequency, the amplitude of the electric field can be obtained directly after the electrode layer, as shown in Fig.2b.

This is the basic concept of an USNN for the tTIS. The neural network has the formula of stimulation, which is modeled from a reference electric field matrix to determine the modulation of beats. By this, the neural network optimizes the electrode currents, which generate a modulation close to the target, from random electrode currents with repeated back propagation. In this method, the approach to find the optimal solution, by gradually changing the currents of the electrodes only, can easily fall into local minima, and hidden layers are added to solve the problem in various ways.

### Loss function

The loss function is expressed as the product of three ratios as follows.

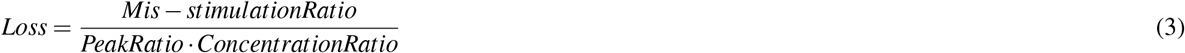

The peak ratio(PR) is the ratio of “the peak modulation of the target” to “the peak modulation of the non-target.” Thus, the largest modulation was applied to the target location.

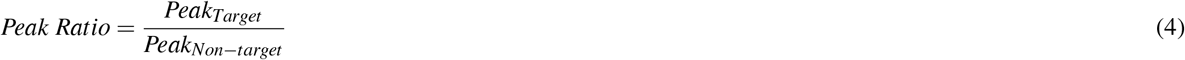

The concentration ratio(CR) is the ratio of “the modulation ratio of the target” to “the area ratio of the target,” so that much of the modulation is concentrated at the target position.

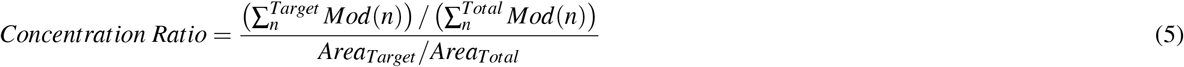

Finally, the mis-stimulation ratio(MR) is the ratio of “mis-stimulated area” to “the target area”. Here, the mis-stimulation is defined as the position where the modulation is larger than the average modulation of the target.

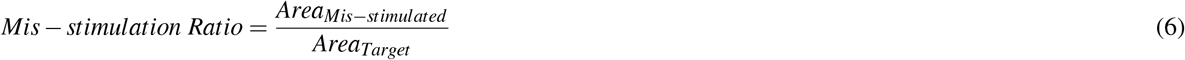

The Adam optimizer^49, 50^, which optimizes the weights of the network, is based on gradient descent. The objective function must be continuous and differentiated at all times. Therefore, the mis-stimulation area was approximated using a sigmoid function centered on the average value^51^.

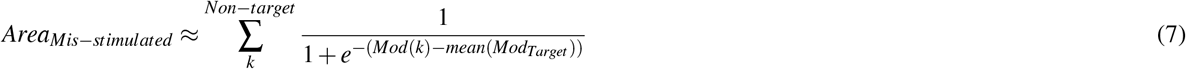

### Detail of computational study

#### Analysis by depth

To verify the performance, the changes in the results according to depth were analyzed. As shown in Fig.3a, by dividing the front and rear end points of the head into nine equal parts on the vertical axis, the position was determined by entering the steps from the front. The horizontal axis is the position translated by dividing the right brain into three from the left, and the height is half of the cerebrum. In this way, the location was moved to a deeper location along the depth index. Each target was 50 nodes of the cortex closest to the location point was selected. All results were compared with the least squares estimation (LSE) results and USNNs results to single frequency tACS.

**Figure 3.**
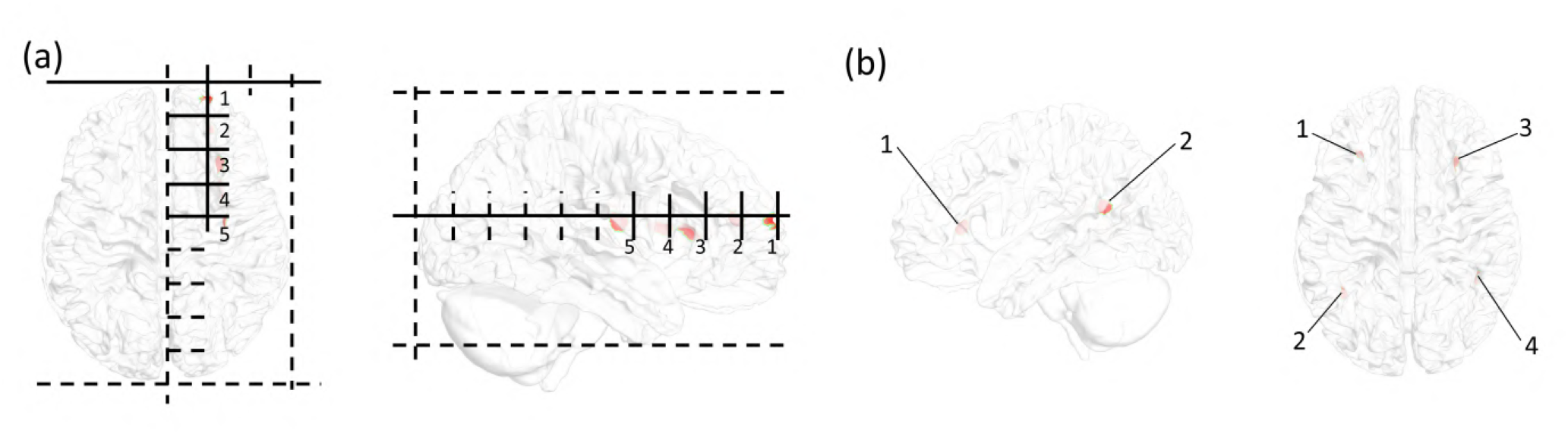
(a) Schematic of the position of the five depth indexes. The shallow part has an index number of 1, and the deepest part has an index number of 5. (b) Index of multi-stimulus position. Position indices 1, 3 in multi-stimulus are the parts of the prefrontal cortex that folds forward, and 2, 4 are the narrow and deep folds on the upper and posterior sides of the brain.

#### Multi-stimulation

The deep region for multi-stimulation was folded forward in the prefrontal cortex (Fig.3b), and deep wrinkles in the vicinity of the intersection of the parietal, temporal, and occipital lobes (Fig.3b). Multi-stimulation was tested for pairs 1-2, 1-3, and 1-4 pairs, and the performance of each part was calculated by dividing the brain into a horizontal or vertical axis. All results were compared with the LSE results and USNN results to single frequency tACS.

#### Comparison with two-pair interfering stimulation

A comparison with the two-pair interfering stimulus, implemented with GA^52–54^ (Supplementary Information), was performed in the insular region^55^. This comparison was carried out for only one head model (ICBM UTHC 1088 766), and the genetic algorithm selected the highest value by performing 100 iterations with 1000 generations.

#### Study environment

All developments and simulations were performed on an Intel i9-9900k, 64GB RAM, NVIDIA Geforce RTX 2070 Super environment. Realistic head modeling and result analysis were performed using MATLAB R2019b, and the finite element analysis of the stimulation was written in Fortran in Visual Studio 2017. The neural network was developed using Tensorflow 2.3.0, and Keras 2.4.0 in Python 3.7.6.

## Results

### Analysis by depth

Figure 4 shows the results for depth, as a surface plot, on a head model, of the optimization using LSE and neural networks for tACS and tTIS. In the case of tACS, the results of neural networks with relatively large stimulation are applied to the target compared with the results of the LSE, but slight oscillation can be confirmed to occur over a wide area. In the case of tTIS, accurate and focused stimulation was performed well, even in deep areas. This is also shown in the box plot of the whole brain model in Fig.5. While the peak ratio and mis-stimulation ratio worsened in the case of tACS according to depth, the case of tTIS maintained a constant level. However, in terms of the concentration ratio, tTIS also worsens, but is always better than tACS. The peak value of the modulation of the target using LSE is lower for all depth indices than others. The two results optimized by neural networks show different superiority depending on the location.

**Figure 4.**
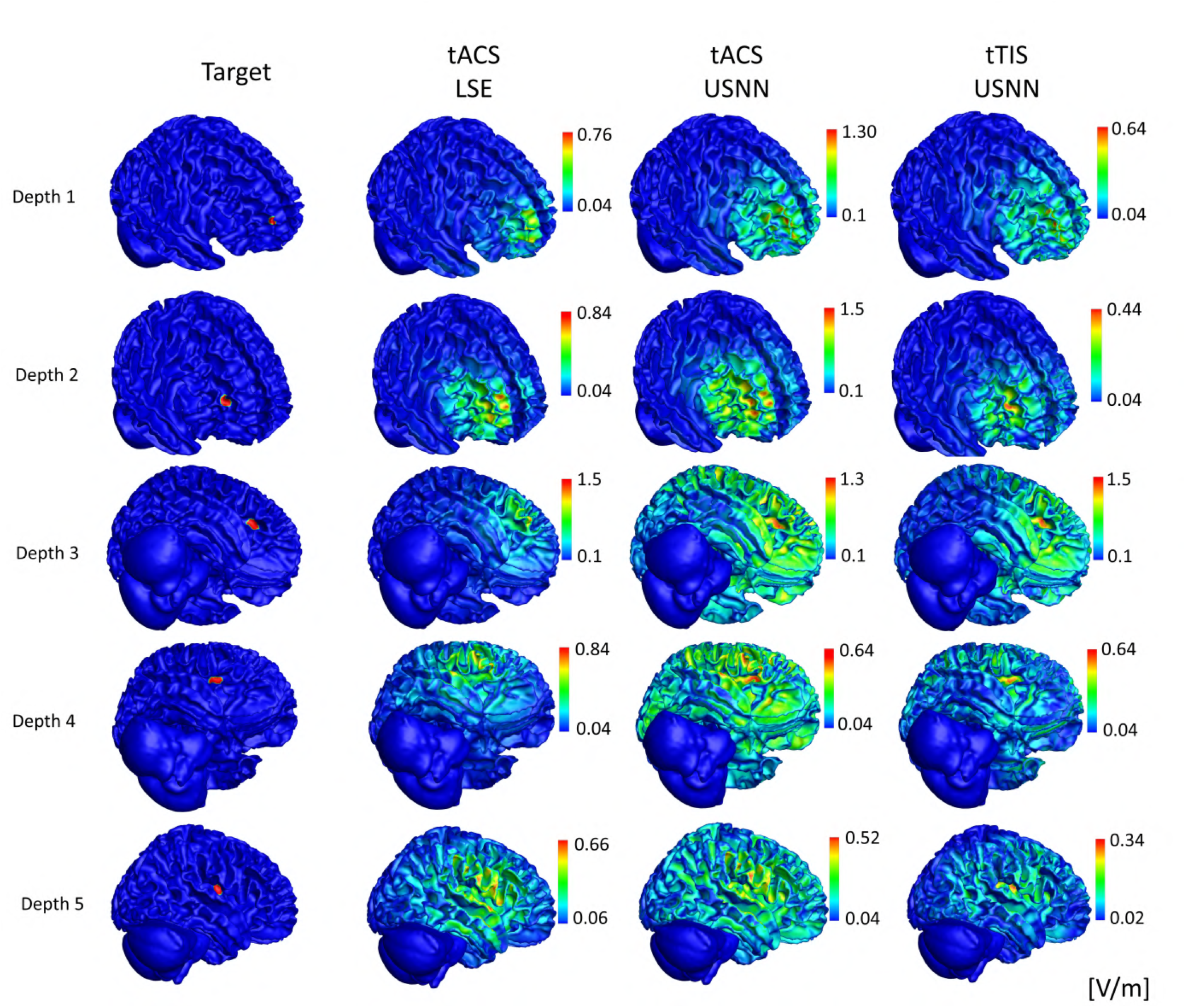
Visual comparison of the modulation generated by the three methods (optimization for tACS by LSE, and for tACS and tTIS by proposed neural networks) according to depth.

**Figure 5.**
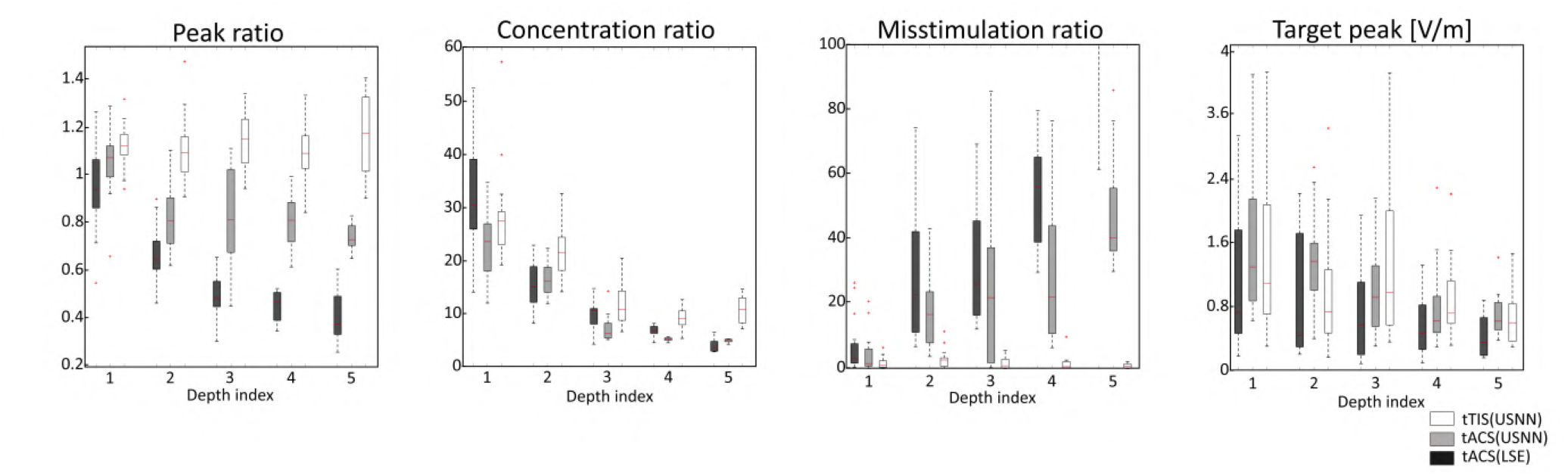
Box plots of 16 head model results for the three methods according to depth about peak ratio, focal ratio, mis-stimulation ratio, and the peak modulation of target.

### Multi-region stimulation

Figure 6 shows the results of optimization of multi-stimulation using LSE and neural networks for tACS and tTIS on a head model as a surface plot. Similar to the test for depth, when optimizing tACS with neural networks, a relatively stronger stimulus than LSE is generated, but weak modulation is formed over a very wide area. This phenomenon is also observed in tTIS but less severe than in tACS. In this head model, tTIS failed to stimulate area No. 4 when multiple regions 1 and 4 were stimulated. Figure 7 shows that a quartile is less than 1 for the peak ratio of area No. 4 on the 1-4 multi-stimulation. Of course, results other than the peak ratio were not good. However, even including this area, tTIS shows much superior accuracy compared with tACS in multi-region stimuli.

**Figure 6.**
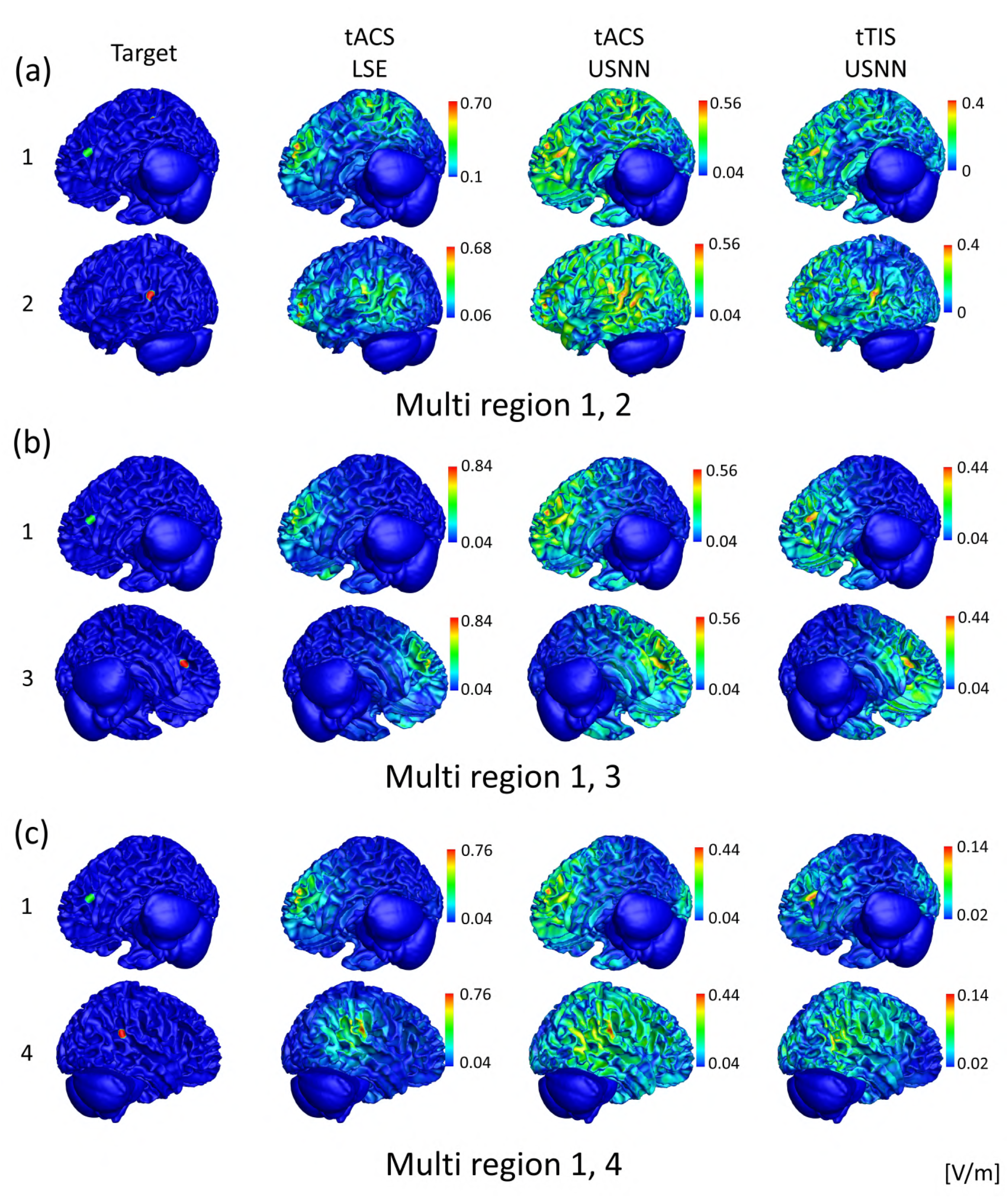
Visual comparison of the modulation generated by the three methods (optimization (a) by LSE for tACS, and by proposed neural networks for (b) tACS and (c) tTIS) used to multi-region stimulation.

**Figure 7.**
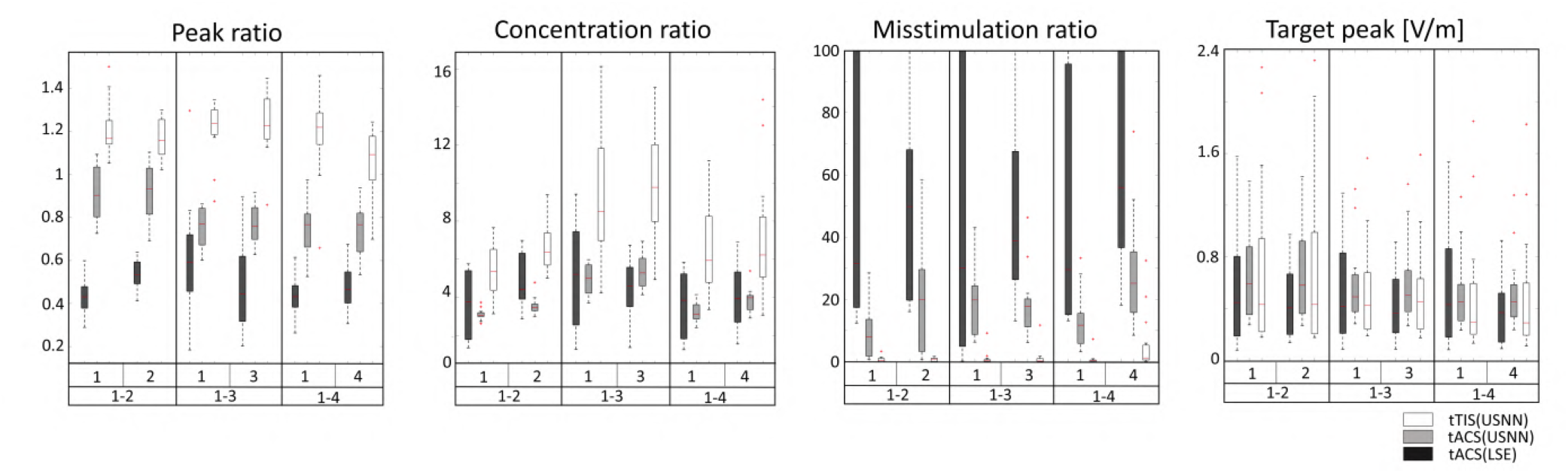
Box plots of 16 head model results for the three methods used to multi-stimulation about peak ratio, focal ratio, mis-stimulation ratio, and the peak modulation of target.

### Comparison with two-pair interfering stimulation

Figure 8 shows the comparison with two-pair and HD-electrode in interfering stimulation results on the insular area in a one-head model. In the case of two-pair electrodes, 17 of 100 GAs had the highest evaluating value. Overall, the case of HD electrode showed remarkably good performance in terms of peak, concentration, and mis-stimulation ratios, but the case of two-pair electrode showed a much higher peak modulation in the target peak

**Figure 8.**
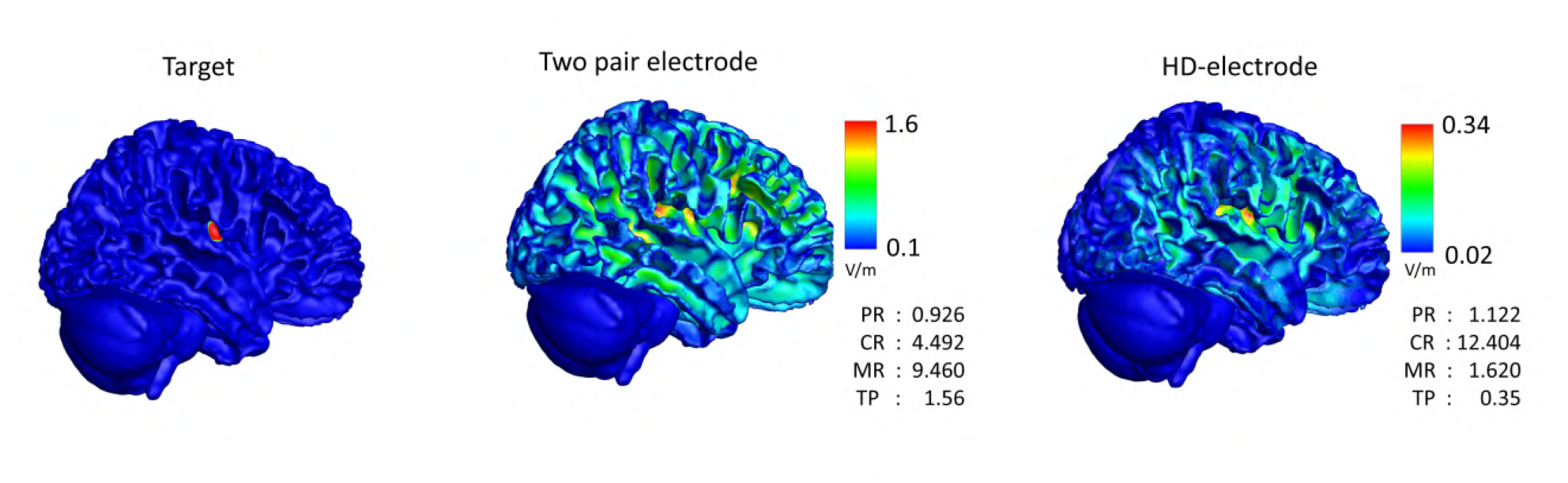
Visual and loss component comparison of the modulation generated by two-pair electrode system and 10-10 electrode system on tTIS.

## Discussion

The performance of the peak ratio and the mis-stimulation ratio of both the LSE and neural network results to single frequency gradually decreased as the depth increased (Fig.5). The result of a single frequency is a linear combination of the results of the reference electric field of each electrode^11, 12^. Because all reference electric fields tend to have a large amplitude in the shallow part near the electrode, the results are naturally poor in the deep region. However, even for a single frequency, when the objective function is used in the proposed method, the performance can be improved to some extent. In the results of tACS using USNN, the concentration was very poor, but relatively strong modulation was generated in the target position. Because of this, the mis-stimulation was significantly reduced compared with the LSE. In the case of an interference stimulus, a very complex form of expression is possible with a nonlinear equation of the reference electric field of each frequency of each electrode as a variable^19, 20^. Because of this, regardless of the depth, the peak ratio almost always exceeds 1 to ensure the strongest stimulus in the target, and the mis-stimulation also maintains a very low level. When the depth increases, there are more non-target regions at a closer distance to the target, so the concentration performance of tTIS also decreases as with single-frequency stimulation. However, it always performs better than the tACS results.

In the case of tTIS, the magnitude of the peak value at the target is disadvantageous for generating large modulation^19^, within constraints where the sum of the injected currents does not exceed 4 mA. The modulation is proportional to the smaller peak values of the two electric fields of each frequency, and they have sources on opposite sides of the target to increase the concentration. However, the results showed no significant difference in the peak value at the target. Moreover, it has larger values in the deep region because of its higher accuracy. Testing different regions showed that both cases have larger single-frequency neural network results in modulation and larger tTIS results in modulation, depending on the location. However, this difference was small. Overall, the disadvantage seems small in modulating the deep region of the brain using interference stimulation in terms of modulation strength.

To date, tES has not been attempted because of poor accuracy at deep depths, but in the case of deep brain stimulation^56, 57^, multiple stimulations have been attempted to stimulate simultaneously two or more deep regions^58–60^. If elaborate stimulation in the deep area is possible due to tTIS and if it can be optimized, tES can also try to stimulate multi-stimulation in deep regions. In Fig.7, the result of optimizing a single frequency stimulation by neural networks is almost always better than the optimization result of LSE. However, generating the strongest stimulation at the target location fails, and some mis-stimulation remains. In the tTIS case, the performance of the peak, concentration, and misstimulate ratios seems better, and the peak values of the target do not appear to differ from the single-frequency results. However, in the results of position 4 of the multi-stimulation 1-4, about a quarter of which failed to stimulate the strongest stimulus to the target and had higher mis-stimulation than the other multi-region stimulation cases in tTIS. For this case, the same result is obtained even if other tests, such as changing the initial value or adding a hidden layer, are attempted. Although a very complex form of stimulation is possible through interference stimulation with a nonlinear equation, the solution of some targets seems to be unable to find in the solution space because of the complex shape of the cerebral cortex. This case was difficult to find when targeting one location. When attempting multi-stimulation, it appeared to have a low probability for a specific combination.

Interference stimulation using different frequencies has been proven to be more effective than single frequency tACS in stimulating the deep part of the brain^20, 21, 26^. In most tTIS studies, a two-pair arrangement, using two pairs of anode and cathode electrodes, has been widely used. Increasing the precision, such as increasing the number of electrodes^26^ or finely adjusting the position^20^, has been attempted, but testing various cases takes a lot of time because the calculation was very complex. A comparison of the results of the optimization using the two-pair electrode arrangement and the results of the proposed method is shown in Fig.8. In the case of two-pair stimulation, it was implemented as a GA, and its solution was determined by testing many times while sufficiently converging without considering the calculation time (Supplementary Information). When the objective function of the GA is implemented as Equation (3), the target is visibly stimulated and is quite concentrated. However, it is not the strongest stimulation in the target region and leaves a certain amount of mis-stimulation. In terms of precision and accuracy, the case in which the current flows from many electrodes at the same time seems much better. However, within the constraints of the electrode current, the peak value of the electric field was greater in the two-pair electrode system. With only two pairs of electrodes, the total current limit of 4 mA can be driven. Thus, HD-tTIS cannot overcome this classical method in terms of strong stimulation.

All tests shown in the results were tested under the same conditions. The same objective function was used in all cases, and the same number of times was repeated with the same hidden layer configuration. The same conditions were implemented; however, this was not the best method. A better result can be obtained by iteratively checking the results and changing the parameters to obtain better results. For example, three ratios of the objective function are multiplied by the same exponent 1 in these results, but the exponent of the focal ratio component can be increased if the concentration is unsatisfactory, to make the response of changing the focal ratio more sensitive. Another example is that the criterion of mis-stimulation can be relatively strict. This example is presented in Supplementary Information. The results according to the architecture of the hidden layer are also presented in Supplementary Information. Estimating electrode currents directly without a hidden layer, such as the method previously used, can easily fall into local minima, and the convergence speed is slow. However, even if only one hidden layer with only two nodes is added, optimization can easily exit the local minima in the solution space. As the number of weights to be optimized increases, the time of one iteration increases, but converges to the optimal value faster and more safely in the appropriately deep hidden layer. Having a wide and fine adjacency complicates the network owing to the high dimension of weights, rather than simply changing the electrode value slightly, so that the network can search for an approach to find an answer faster.

Although simulations enable precise optimization in a short time, this research inherently has limitations in computational studies. The electrode system is ideally designed, and in practice, precise implementation of the attachment and control of many electrodes is difficult. The difference between the simplified head model and real head can cause errors in the precise simulation results. Nevertheless, this method can help to approximate the optimal electrode currents in other feasible electrode systems. In future studies, validation of tTIS implementation will be necessary in practice.

## Conclusion

We proposed a method to optimize the tTIS of a high-resolution electrode system based on USNNs, and we analyzed the possibilities and performance from various cases. The proposed method shows very precise and accurate performance in the tTIS optimization problem, which remains a difficult problem. The method also shows that performing fast calculations enough to apply is possible. As the problem of modulating oscillations in the deep brain becomes more accurate, it is expected to be the basis for more clinical testing and verification in the field of brain stimulation using electric fields.

## Acknowledgements

This research was supported by the KBRI Basic Research Program through the Korea Brain Research Institute funded by the Ministry of Science and ICT (21-BR-01-13 and 21-BR-03-01) and by the National Research Foundation of Korea funded by the Korean Government under grant (NRF-2019R1A2C1011270).

## Author contributions statement

S.B. and C.L. conceived the initial idea. S.B. conducted all optimization and numerical simulation. S.B. and B.K, designed the neural network. C.L. created the models and the software for finite element method. All authors revised and approved the final manuscript.

