## supplementary for "A computational study on the optimization of transcranial temporal interfering stimulation with high-definition electrode using unsupervised neural network"

### Supplementary Information

#### Genetic algorithm for two-pair tTIS

The electrode value optimization of the two-pair interference stimulus was performed using a genetic algorithm. The composition of the chromosome consists of six terms as (electrode position by frequency) and (electrode intensity by frequency), as shown in Fig. S1. The electrode position has an integer value of 1 to 69. The electrode strength has a real value of -1 to 1 and is then normalized in order for the sum of the injected currents to be 4 mA.

The initial population starts with 1000 randomly generated individuals, preserving 100 elites per generation. Here, the evaluation value uses the same function as the loss function of the proposed neural network. Based on the rank, 500 parent 1 and parent 2 are selected, and crossover is carried out. The crossover generates two offspring that inherit the items of parent 1 and parent 2 randomly for each chromosome position. Rank-based selection and random position crossover are intended to slow the convergence speed so that it can converge to the correct answer slowly, avoiding the local minima as much as possible without being constrained by the calculation time. The mutation changes the positions corresponding to 20% of all chromosome positions of the crossed children to random values. When these 1000 generations were repeated 100 times, 17 results converged to the best value among the test results.

#### Performance according to the loss function

The loss function of the main test used in this study is expressed as the product of the peak, focal, and mis-stimulation ratios. The results of using only each component as a loss function are shown in Fig.S2a-c.

In the case of the peak ratio, because it is a value of one position representing a divided region, it cannot always guarantee a continuous and differentiable solution space. The peak ratio has reached a fairly high place, but the peak ratio is rather low compared to when the mis-stimulation is a loss function, and it can no longer go to a high place. This component only guides the result so that the target can be stimulated the most strongly in the early or final fine-tuning stage. Nevertheless, when only the peak ratio is used as the loss function, the peak ratio has a high value. However, large and small mis-stimuli spread throughout the brain.

When only the mis-stimulation ratio is used as the loss function, a relatively strong modulation is generated at the target position, and all modulations elsewhere are smaller than the average of the target. However, weak modulation is spread throughout the brain, and the peak value of the target is low compared with other results. When only the focal ratio is used as the loss function, the entire modulation generated is concentrated near the target. However, the results showed that the other areas next to the target were stimulated more strongly than the target.

When using the product of the three items, each item yields slightly worse, but overall compromises. The strongest stimulus was generated in the target, the overall modulation was relatively concentrated toward the target area, and the mis-stimulation was smaller than the area of the target. In addition, the target peak value was higher than the other results because avoiding mis-stimulation is possible while being focused.

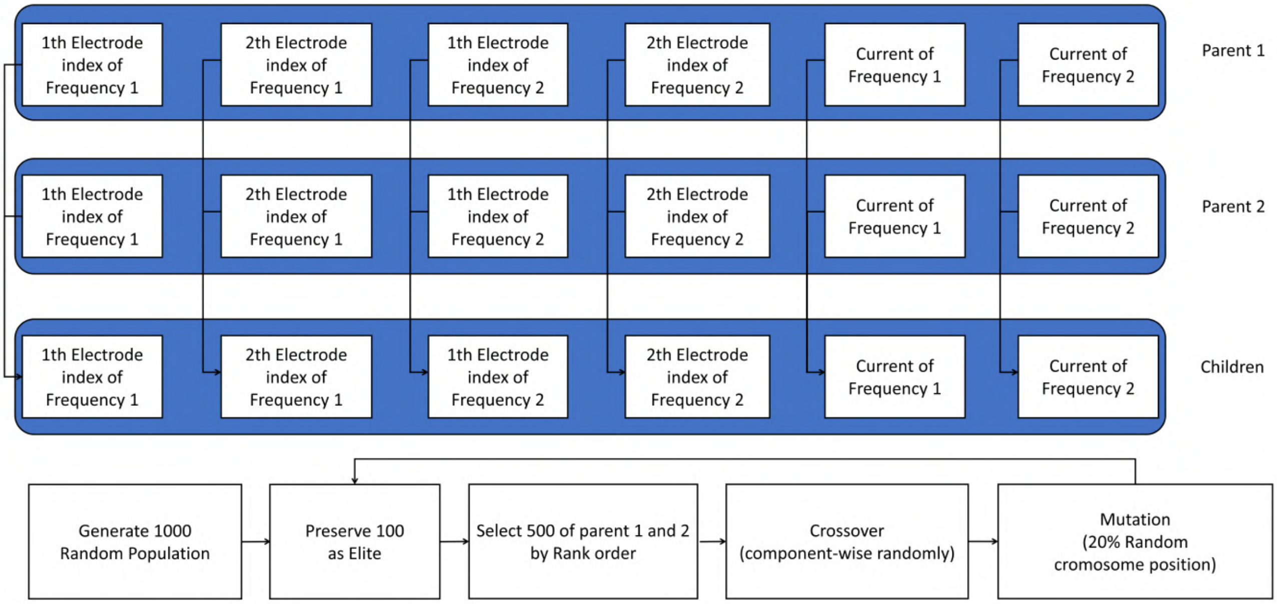


**Figure S1.** Chromosome flow chart of the genetic algorithm for two-pair interference stimulation.

After checking the results, it can be adjusted according to the purpose by modifying the loss function. A simple example is presented in Fig.S2. Because the concentration of the result of producing three items equally was disappointing, the exponent of the focal ratio was set to 3 to make the reaction very sensitive to the concentration ratio. In addition, to reduce the size of the part that is mis-stimulated on the side of the target, the criteria for mis-stimulation were lowered by half. The results showed that modulation was more concentrated than the previous results, and the size of the lateral mis-stimulation was reduced.

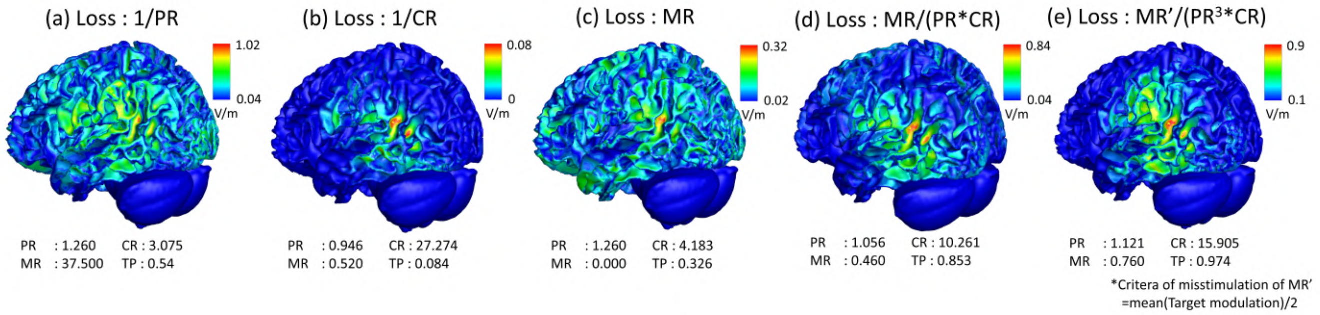


**Figure S2.** Optimization results according to the loss functions.

### Performance according to hidden layers

If the electrode currents are directly determined using the gradient descent method without using a hidden layer, the solution easily falls into the local minima. However, by adding a hidden layer of Relu consisting of only two nodes, the network can avoid these local minima and find a better solution. As in the architecture used in the main results of this study, if five layers are configured with an exponentially increasing number of nodes, such as 2, 4, 8, 16, 32, it converges to a better result much faster.

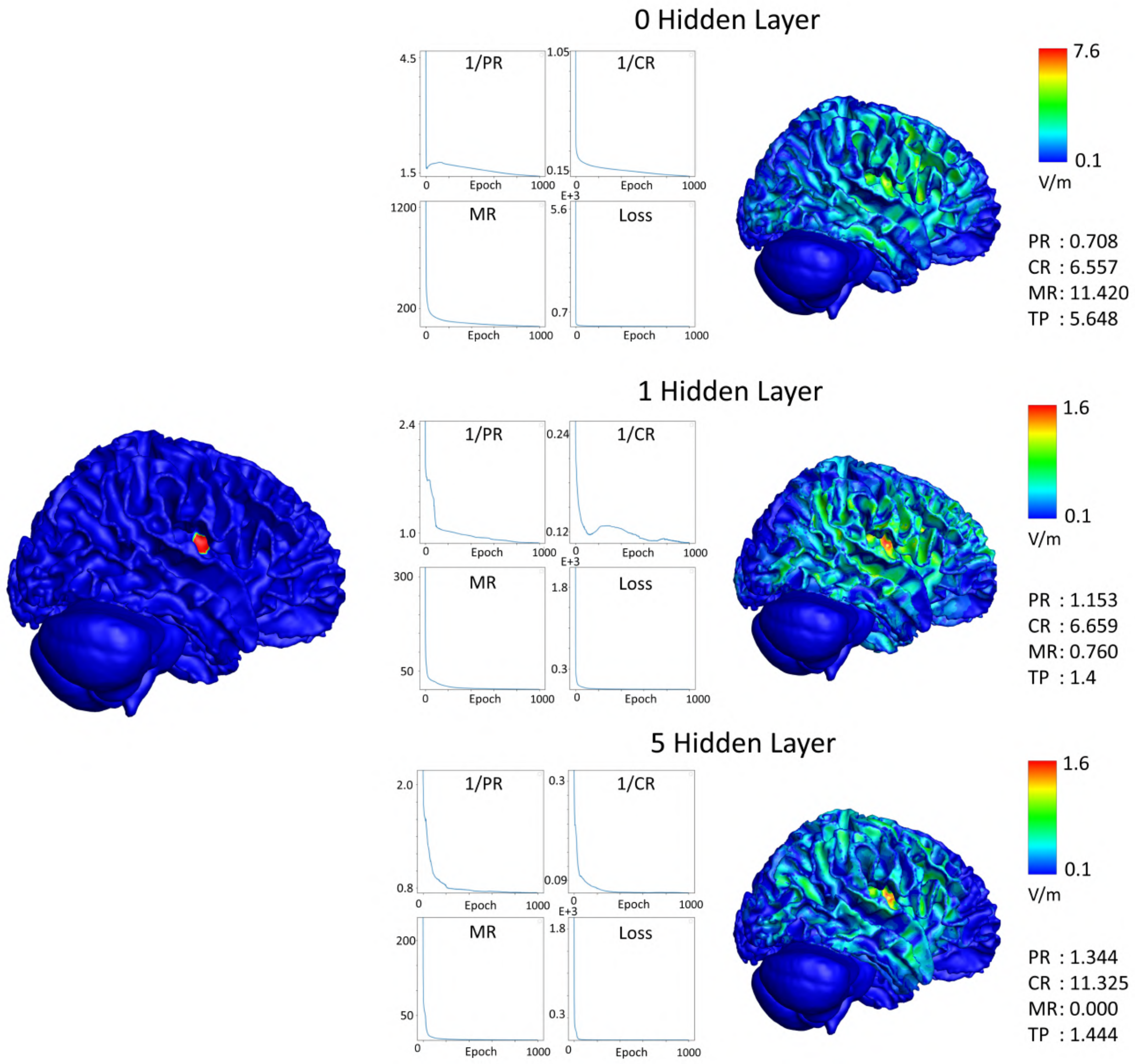


**Figure S3.** Optimization results according to change in hidden layers. The graph on the left shows 1/peak ratio, 1/focal ratio, mis-stimulation ratio, and the product of three items for the epoch. When only one hidden layer is used, it is composed of 2 nodes, and when using 5 hidden layers, 2, 4, 8, 16, 32 nodes are configured to increase exponentially.
